# What “unexplored” means: Mapping undersampled regions in natural history collections

**DOI:** 10.1101/2024.02.09.579602

**Authors:** Laymon Ball, Ana M. Bedoya, Sheila Rodriguez Machado, Diego Paredes-Burneo, Samantha Rutledge, David Boyd, David Vander Pluym, Spenser Babb-Biernacki, Austin S. Chipps, Rafet C. Ozturk, Yahya Terzi, Prosanta Chakrabarty

## Abstract

We examined global records of accessible natural history voucher collections (with publicly available data) for terrestrial and freshwater vascular plants, fungi, freshwater fishes, birds, mammals, and herpetofauna (amphibians and reptiles) and highlight areas of the world that would be considered undersampled and sometimes called “unexplored” (*i*.*e*., have relatively low, or no evidence of, past sampling efforts) under typical Western-scientific descriptions. We also question what “unexplored” may actually mean in these contexts and explain how retiring the term in favor of more nuanced phrasing can mitigate future misunderstandings of natural history science.

## Introduction

People have been collecting natural history specimens for centuries and these specimens have been used for gathering valuable data about our changing planet [1–3]. While the number of collections from some regions has steadily increased over time, many other regions have persistent gaps in their collection records. It may be that no naturalist history collectors have ventured into that space because of inaccessibility (*e*.*g*., the tops of a tepui, the bottom of a tsingy), political reasons (*e*.*g*., it is protected indigenous land, or in a war-torn/unsafe zone), or sometimes these areas may have simply been overlooked [4–5]— for example, some areas may be presumed to be biodiversity-poor (e.g., deserts) and overlooked in favor of well-known areas of high species richness (e.g., coral reefs, rainforests). Alternatively, it could be that specimens from these areas actually exist but have not been digitized— digitization being the “translating [of] metadata associated with a physical specimen object into flexible digital data formats” [6]; funding is a known barrier to digitization at institutions globally [7]. Or collections from these areas may exist, but they remain in a “blind spot” because they have not been aggregated into publicly accessible global databases such as the Global Biodiversity Information Facility (GBIF) which is the single largest biodiversity repository for such data. In this paper, we aim to shed light on what could be meant by “unexplored” by examining the different factors that explain the dearth of apparent collection activities in some parts of the globe.

Collections-based natural history research is the work of observing, securing, obtaining, and preserving wild organisms for future study. Data collection from these organisms can include photographing, measuring, and obtaining blood or tissue samples for future molecular and/or biochemical analyses. These organisms are preserved in a fixative (such as formalin), dried, skinned, and stuffed, or otherwise prepared for long-term comparative use as a reference “voucher” specimen [8–10]. Vouchers are evidence of the existence of these organisms in a particular place and time (that can be compared with specimens from other time periods and locations). These reference specimens document morphological, genetic, and phenological variation of a species or population and are used as proof of a species new to science (as type specimens). Scientific specimens can be used for countless research applications including as part of the “extended specimen” concept [11–14]. In particular, digitized collections facilitate large-scale studies on critical questions in global change biology [6,15], and GBIF-enabled research extends across all major science disciplines [16]. Publicly available natural history collections also facilitate opportunities for international and interdisciplinary collaboration [12].

Unfortunately, natural history collections are in serious decline [17]. Part of this decline is the shifting in focus of traditional natural history museums away from collections research towards a focus on public exhibits and sustainability [18], as well as cultural and political resistance to the idea of collecting certain organisms [19, 20]. It is also related to a decline in basic science research funding in natural history and the growth of more applied scientific research with economic goals [18], leading to greater inequalities and mistrust between researchers from resource-rich versus resource-poor countries or institutions (see “Institutions Holding Collections” table from [5]). These inequalities lead to the pressing need for institutions conducting collections-based research to connect and share opportunities more equitably.

Here, we aim to map the regions of the globe highlighting areas where there are relatively few or no publicly available natural history records and consider the potential reasons behind these deficiencies. We focus on terrestrial and freshwater habitats, and exclude the vast and still relatively poorly explored and enormous marine realm [21–23]. Thus, we limited our searches to terrestrial and freshwater vascular plants, and to terrestrial vertebrates (i.e., birds, herpetofauna, and mammals) and freshwater fishes. Though oceans comprise the most abundant habitat on Earth, most natural history research has focused on terrestrial environments [24, 25]. Therefore, the lack of collections in these terrestrial areas may be more notable and informative than that of the relatively overlooked and vast oceans.

Regional gaps in specimen collection may not be consistent across taxonomic groups [4]. For example, an area depauperate in collections of mammals may not be so for freshwater fishes, and vice versa. Myer [26] identified both taxonomic and geographic biases in plant collections, particularly towards North America, Western Europe, and Australia, and found that outside of these regions, high collection coverage tended to be associated with a specific botanical program, such as the Missouri Botanical Garden’s multi-decadal efforts in Madagascar [27]. Therefore, we attempt here to address regional collection biases for each taxon examined separately and identify “blind spot” regions that lack scientific collections for multiple taxa. We highlight areas historically deemed “unexplored” in order to better recognize the reasons behind sampling biases and raise awareness about habitats that might be worthy of future natural history research and preservation. Future research may include aiding smaller, private, or non-digitized collections to become part of the global collections infrastructure by helping to make their vouchers and associated data accessible and public. We also aim to identify areas needing additional conservation protection if the current status is insufficient. Our work may help recent advances in predictive modeling that can be used to track regions with a high potential to hold hidden biodiversity [28]. We hope this study will help identify the gaps in our understanding of the world’s biodiversity and draw attention to how we identify areas that have been potentially overlooked by those interested in the exploration and conservation of the natural world.

## Results and Discussion

We examined over 55 million (55,200,291) occurrence records from each continental region (Table 1) as defined herein, and compared the patterns of sampling efforts for natural history collections for all six major taxonomic groups. We focus on the shared poorly-collected regions and those unique to each of the six taxonomic groups included. In some countries, an artifact of older digitized data is that localities may be listed as simply “Brazil” or “Peru,” causing the centroid (relative midpoint) of that country to be the area highlighted as most “collected.” We avoid descriptions of such artifacts here. However, we note clear cases where the presence of a local museum, herbarium or fungarium increases local specimen collections relative to other regions.

**Table 1:** Specimen records by continental region and taxon.

|  | South America | North and Middle America | Africa | Australia and Oceania | Eurasia | Antarctica |
| --- | --- | --- | --- | --- | --- | --- |
| **Birds** | 361,711 | 1,856,474 | 227,471 | 445,378 | 535,312 | 2,202 |
| **Fishes** | 360,685 | 1,484,529 | 188,976 | 179,446 | 443,824 | 1,128 |
| **Fungi** | 162,228 | 1,280,467 | 59,101 | 416,647 | 1,972,661 | 9,967 |
| **Herps** | 436,020 | 2,278,628 | 256,039 | 825,493 | 251,687 | - |
| **Mammals** | 279,689 | 2,033,383 | 317,256 | 278,725 | 555,010 | - |
| **Plants** | 6,723,374 | 10,271,258 | 2,089,744 | 6,342,071 | 12,269,675 | 4,032 |

Data, originally obtained from GBIF, were filtered to exclude observational data without vouchers and to only include specimen records with geographic coordinates in terrestrial systems, see “Materials and Methods” section for more information. Note: “Herps” is a shorthand referring to “Herpetofauna” i.e., amphibians and reptiles.

### South America

Collecting records across South America show that across taxa, publicly available collections are more numerous in mountainous regions and well-known biodiversity hotspots (*i*.*e*. the Andes, Atlantic forest, and the Guiana Highlands; Fig 1). This pattern is likely a result of collecting efforts being centered in well-known areas of high species richness. Similarly, digitized collections along the Andean mountains show a latitudinal gradient in sampling effort. This pattern is consistent across taxonomic groups and is likely linked to a well-documented latitudinal biodiversity gradient [29]. Every group examined appears to have a high number of collections in the Atlantic Forest of Brazil and Uruguay relative to dryer adjacent regions including the savannas of the Cerrado. However, estimates of undersampled terrestrial vertebrate diversity made using publicly available data have identified hotspots like the Andean mountains as among the most undersampled regions [4]. Assessing the extent of the scarcity of data collection in highly biodiverse regions remains a challenge. There are relevant records stored in private collections, as well as specimens at local institutions which have not, or have only partially been aggregated into digital repositories (e.g., Museum of Zoology of the University of São Paulo, Cartagena Botanical Garden “Guillermo Piñeres”, the National Museum of Brazil, the *Museo de Historia Natural* in Lima), which remain hidden to the international scientific community.

**Fig 1.**
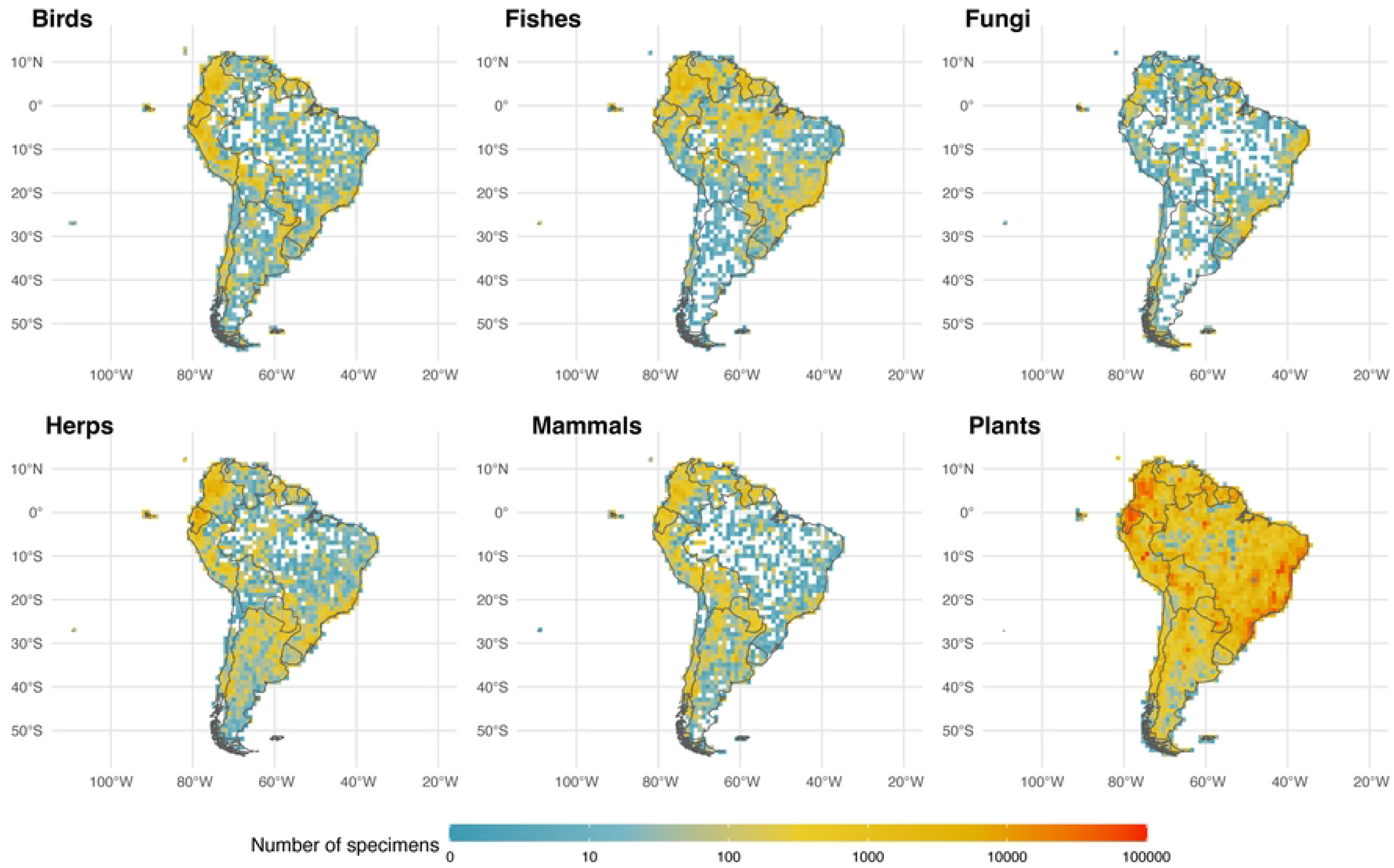
Maps of collecting efforts on South America. Sample (voucher) quantity for each taxon considered in this study is shown. Red colors represent 100K+ specimens, white and very light colors represent the so-called ‘unexplored’ areas or areas with few collections.

The Amazon basin, one of the largest and most biodiverse regions of the world (holding 10% of all named plant and vertebrate species; [30–35]), has lower sampling across taxa (particularly for fungi and mammals; Fig 1). Large sections of the Amazon remain largely undersampled, with botanical census data covering <0.0003% of the region [32]. Other lowland areas with fewer publicly available records across taxa include the Orinoco basin, the Atacama Desert, the southern portion of the Chaco, and the Patagonian grasslands. While fewer digitized collections in the Atacama may be representative of the region’s low diversity for the six taxonomic groups examined here, accessibility and armed conflict may be relevant factors limiting exploration in the other regions. Scarcity of roads or navigable water bodies are a challenge to transportation to and from large sections of those regions, increasing the costs, risks, and logistics for field exploration [36–37]. Political and civil unrest in Latin America have historically limited collecting in these and other regions and may have kept many collections there from being part of a global digitized database. This lack of digitization may be reflected in the dearth of collections represented in what are very biodiverse countries. In fact, such biodiversity is related in part to a positive relationship between forest cover and the intensity of armed conflict [38–41].

Specimen data for plants are the most abundant compared to the other taxonomic groups examined here. However, it has been estimated that there are zero publicly available botanical records from about 10% of tropical South America [32]. A higher number of botanical collections may be due to higher plant species richness compared to other groups. In addition, the common practice of collecting multiple replicates per specimen to be shared across herbaria could result in duplicated digitized records and in a higher probability of a given specimen being digitized.

Freshwater fishes are the most speciose group of vertebrates that we examined, and their species richness peaks in South America, specifically along the main tributaries of the Amazon River basin (circa 7000 species; [42]). The Amazon Basin and major tributaries appear much better collected (and digitized) than the Orinoco River Basin which covers much of Venezuela and Colombia. In Venezuela, only the Universidad Central has published data in GBIF (for insects), and many are privately held. The “La Plata” region (including the Parana and Paraguay) in east central South America is also poorly sampled and/or digitized relative to the Amazon. Notably, the Deseado River estuary, Lake Musters, and Nuevo Gulf in Argentina are well-sampled for fishes. Compared to the other groups examined, there are comparatively fewer records of fungi. Intensively “explored” regions for this group may be a consequence of the presence of institutions where this taxonomic group has been well sampled and where collections have been digitized. For instance, southeastern Brazil appears more intensively sampled because the Herbarium at Federal University of Pernambuco is located in this area and has the largest collection of fungi in Brazil [43]. Relative to northern South America, Brazil appears sparsely sampled for specimens of mammals, herps, and birds, especially relative to plants and fishes.

### North and Middle America [including landmasses in the Gulf of Mexico, Caribbean and West to Hawaii]

North America is one of the most consistently sampled continents with all taxonomic groups exhibiting similar decreases in sampling effort along a south-to-north latitudinal gradient and few areas devoid of specimen samples of any group south of Canada (Fig 2). Excluding Greenland, interior Canada is the largest gap for all plant and animal groups examined, from Ontario northwest to NW Territories and east to Newfoundland and Labrador. Canada may remain relatively undersampled in part because of its enormous size; the lack of knowledge about its abundant freshwater resources has been previously noted [44]. Relative to adjacent countries it appears Nicaragua is poorly sampled for all vertebrate groups and fungi despite having an abundance of biodiversity-rich regions including the Mosquito Coast, Lake Nicaragua, and the largest tropical rainforest north of the Amazon [45]. Otherwise, most of Central America and the Caribbean/Greater Antilles appear well sampled with a clear gap for Cuba, due more so to conflicting politics and policies (at least with the United States) than to a lack of biodiversity [46]. Conservation work in Cuba is among the best in the region [47] and the Museo Nacional de Historia Natural de Cuba in Havana has digitized collections but these are not publicly available. Perhaps not surprisingly the United States is very well sampled, and climatic differences between the wetter, more humid east versus the more arid west can be seen when comparing fungi and fishes from those regions; with the more temperate Pacific Northwest being an exception.

**Fig 2.**
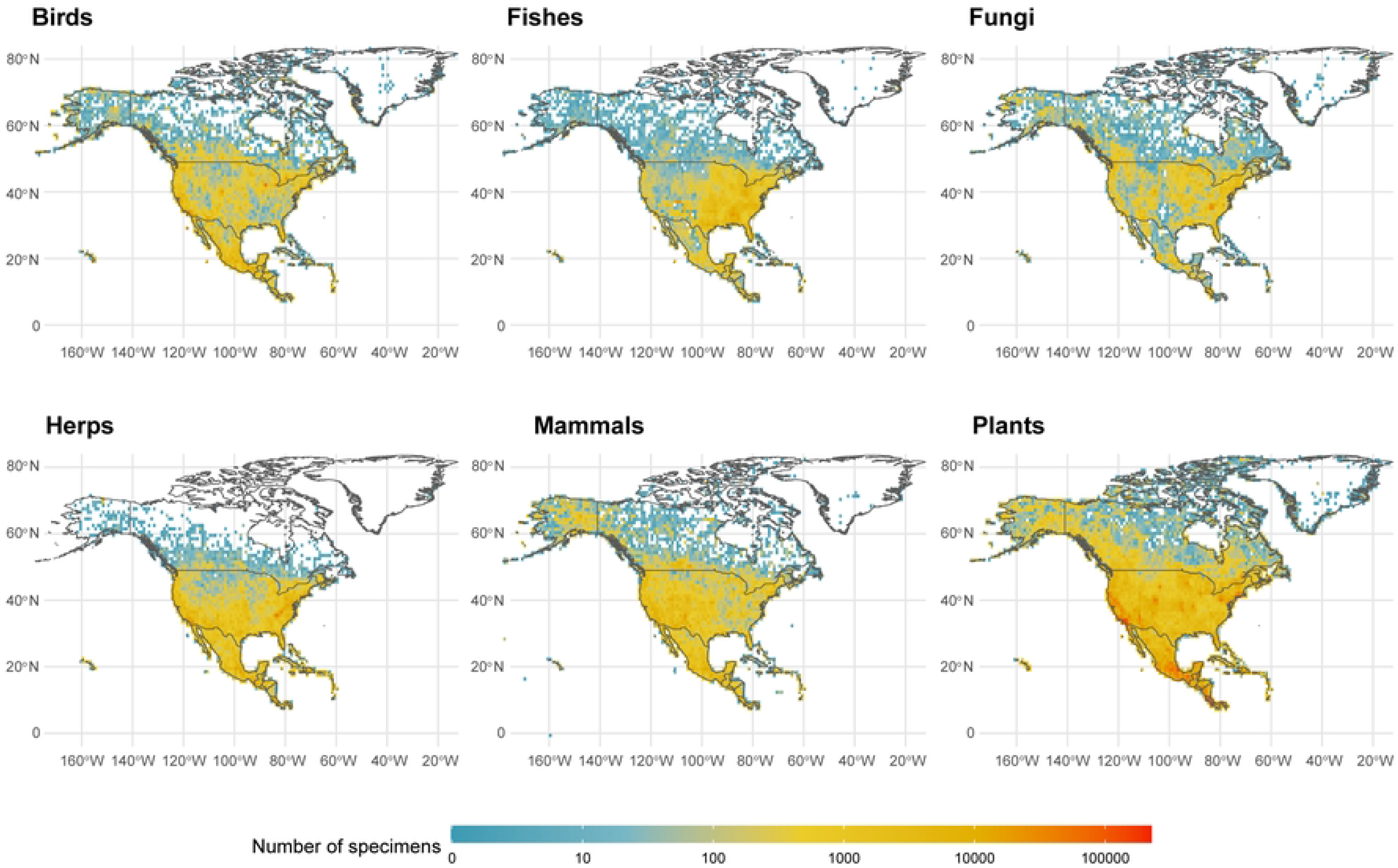
Maps of collecting efforts on North America and adjacent regions. Sample (voucher) quantity for each taxon considered in this study is shown.

Mammals have weaker collection records from the southeast and midwest United States than other vertebrate groups. Specimens of amphibians and reptiles (herpetofauna) display the sharpest decline moving northward of all examined groups, with few specimens collected in and east of the Rocky Mountains from Montana to Illinois. Fishes are, unsurprisingly, sparsely collected from the northern Rocky Mountains south to central Mexico including adjacent desert regions. Although most vertebrate groups display strong holdings from the southeastern United States, non-coastal southern Appalachia is relatively poorly represented in Bird collections, as are Montana, Wyoming, and Utah in the west. Plants are the most consistently sampled group across North America with few, if any, undersampled regions, while Fungi have the highest number of unsampled areas, especially in and around the Chihuahuan Desert.

### Africa [including Madagascar and adjacent landmasses]

Perhaps more than any other region, Africa’s “unexplored” areas likely reflect the sparsity of digitization of regional collections as well as the general lack of collections (Fig 3). A wide variation in climate such as the dry Sahara region versus sub-Saharan tropical regions are also important explanations for variation in collecting effort within the continent. A striking juxtaposition is seen between relatively well-sampled Madagascar and adjacent islands such as the Seychelles of the southeastern coastline, particularly for plants, which is likely a result of sampling efforts from the Missouri Botanical Garden which has a permanent base situated on the large island, and a long history of botanical exploration there; today, the Garden accounts for an estimated 72% of vascular plant collections from Madagascar. Unfortunately, this means the largest collection of Malagasy plants exists well outside of the island which would hinder the growth of local knowledge on these organisms.

**Fig 3.**
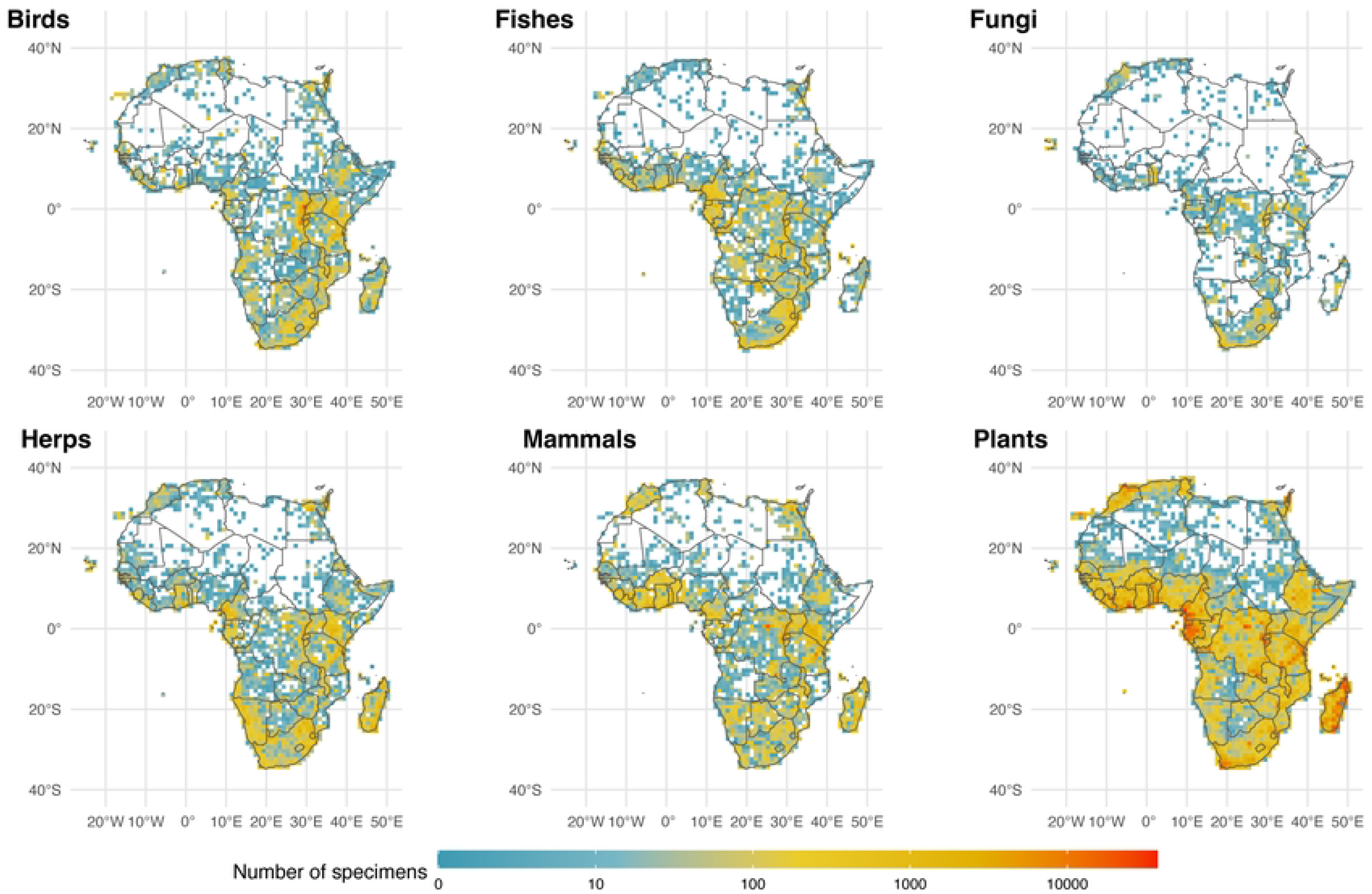
Maps of collecting efforts in Africa and adjacent regions. Sample (voucher) quantity for each taxon considered in this study is shown.

Herps show a similar pattern of natural history sampling effort to plants with a diagonal band across the tropical southern equatorial regions. Collections for most groups are generally tracking with humidity and rainfall, save for fungal and fish diversity which is likely higher than represented by the few collections that have been made across the continent, including Madagascar. Notably, Cape Verde is a bright spot for fungal collections on the West Coast of Africa almost entirely accounted for by collections from lichenologists at the Senckenberg Research Institute and Natural History Museum in Frankfurt, Germany, demonstrating the outsized impact a single research program can have on perceptions of whether an area has been ‘explored’ [48]. The Great Rift Valley region is particularly well sampled for mammals, as are the southern reaches of West Africa (from Liberia to Benin). Madagascar, given its high proportion of endemic mammal species, has few museum specimens; this is likely due to the conservation status of many of the species, including the endangered endemic lemurs.

Despite being largely tropical, Nigeria appears to be a surprising gap for several groups relative to adjacent countries. Similarly, Somalia, the Central African Republic, the Democratic Republic of the Congo, and Angola are gaps for several groups. The sparsity of collections in these regions can be attributed to a combination of complex factors including a history of colonization, lack of scientific infrastructure and institutional support, geographic accessibility, and historical and present-day political instability (particularly in central Africa; plant collections have been noted to decline during periods of war [49]). While not exclusive to these poorly collected regions, cultural factors are also an important consideration (e.g., in some areas there may be cultural beliefs that discourage the collection of natural history specimens; [34, 49]). We call for collections in Africa to join the GBIF network but also call on institutions outside of Africa with African collections to publish their records (some Western museums have collections from the region that are not yet digitized). Egyptian collections from Cairo, Alexandria, and along the Red Sea for several groups are notable and likely were a target for several important historical collections. Interior reaches of Africa, outside of the desert regions, including the Okavango Delta still have relatively few collections. However, given that regions celebrated conservation status, efforts to collect in this area should be well-regulated. Notably many collections from this region are in Europe and the United States, and efforts to digitize existing collections and create new regional collections should be considered as they would benefit local knowledge.

### Australia and Oceania

Unsurprisingly, the large swath of arid land through central Australia is less densely sampled than tropical northern regions and the temperate south, due to the relative dearth of species biodiversity in this arid climate, particularly for large vertebrates (Fig 4; [50]). This portion also remains logistically difficult to access for collecting, due to its desert climate and relative lack of travel infrastructure (i.e., roads). Tasmania appears to be a hotspot for specimens of all taxonomic groups, as does the region near Perth in the Southwest, due in part to their more tropical climates, easier accessibility, and the presence of natural history museums in each locale [43, 50].

**Fig 4.**
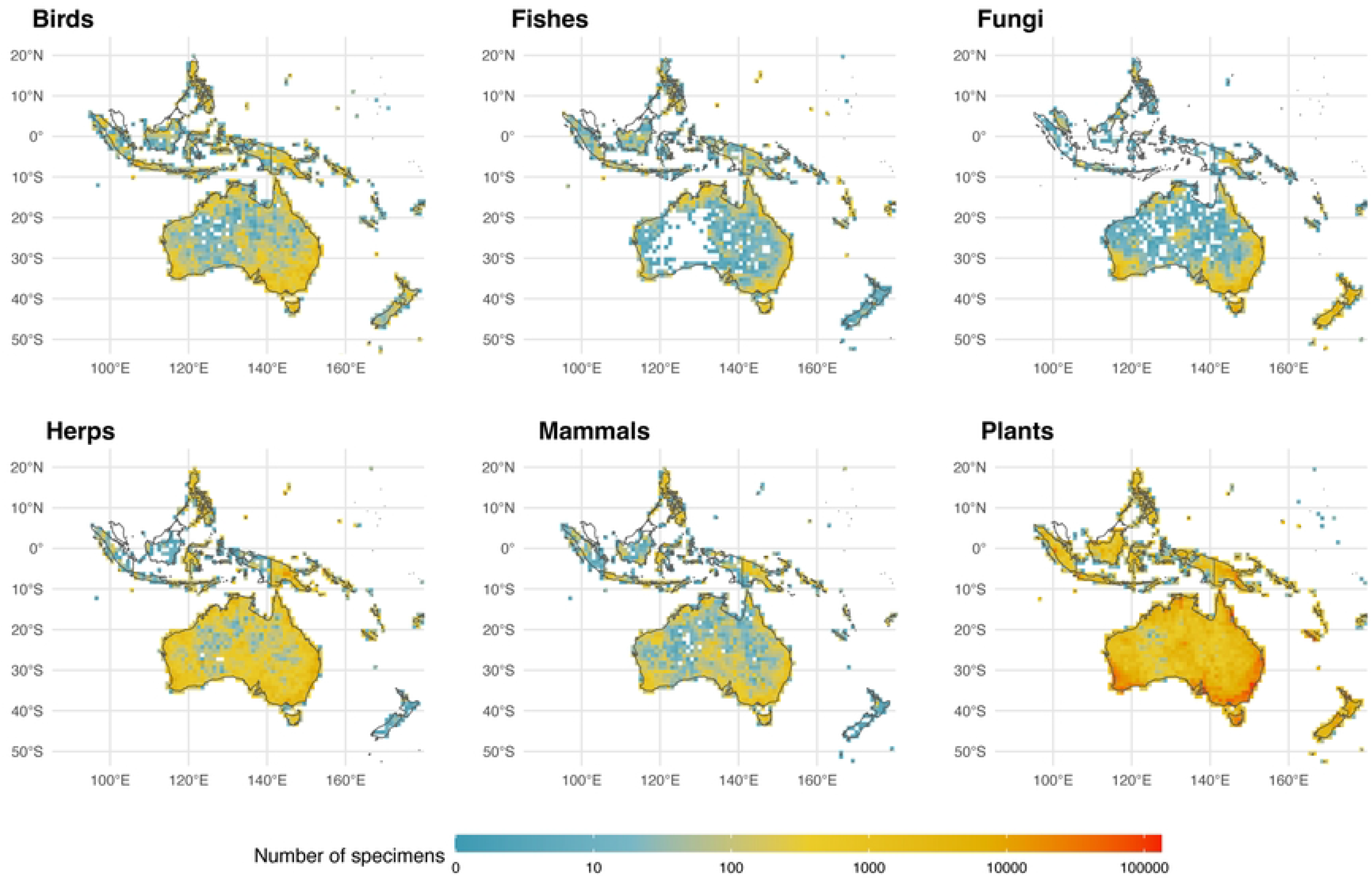
Maps of collecting efforts on Australia and adjacent regions in Oceania. (such as the Indonesian archipelago and New Zealand). Sample (voucher) quantity for each taxon considered in this study is shown.

A sharp divide exists in New Guinea between New Guinea (the western Indonesian side often called “Irian Jaya”) and the Eastern side of the island, Papua New Guinea, which has been a target for many natural history expeditions [51-53]. Another noticeable division occurs between north and south of the New Guinea Highlands, with the south being poorly collected versus its northern counterpart. This is the case for both the Indonesian and Papuan halves of New Guinea, most likely due to the heavy monsoon flooding that seasonally submerges areas in that portion of the island [54].

Despite its tropical climate and abundance of biodiversity, there appear to be few collections from Indonesia, particularly for fungi. Contrastingly, the Philippines (central top of map) are densely collected relative to Indonesia, again showing how Indonesia appears poorly collected for natural history specimens. It is also possible that Indonesia only appears relatively “unexplored” because of a dearth of accessible digitized specimens (likely in collections in the West). Indonesia has been a target for museum-based collections since occupation by the Dutch and Wallace’s early expeditions to the ‘Malay Archipelago.’ Few if any of the specimens from southeast Asia collected by Wallace or used as type material by Western zoologists are curated by local institutions. Specimens collected from this region more recently are curated in large institutions like the Museum Zoologicum Bogoriense in Java, Indonesia, which has yet to have its extensive collections digitized. Notably, no museum collection from Oceania is listed among the 73 of the world’s largest natural history museums and herbaria from 28 countries with the nearest of these surveyed collections being in India and Australia [5]). Representation of people from this region and digitization of local collections should be a priority before labeling this entire archipelago “unexplored.” The absence of a fungarium (despite there being herbaria) is an oversight and can be linked to why there are few fungal collections in an area that should be a hotspot for mycologists. Borneo appears remarkably underrepresented in natural history collections for herps and mammals, particularly relative to nearby Java; the lack of mammal and herp collections from New Zealand is less surprising because of the relatively few species of these groups known to exist there. Mammals and herps are particularly well collected in the Philippines and Papua New Guinea making adjacent areas even more glaringly under-sampled. Birds and plants appear to be the best sampled taxa from Indonesia, perhaps as a result of historic collecting efforts but large gaps remain in New Guinea, Borneo, Sulawesi, and Sumatra.

### Eurasia

Much of Europe has been well collected and the specimens digitized from the region are among the highest on the planet; in stark contrast to the rest of Eurasia except the far East (particularly Japan and Taiwan which have substantial collections and active natural history researchers; Fig 5). The relative absence of digitized collections from India for plants, fungi and mammals is unexpected, given the region’s rich biodiversity. Sri Lanka appears well-sampled relative to India for several groups which may reflect a greater accessibility to collections and digitization of specimens from that island nation. South Korea, Nepal, and Taiwan appear to have a high number of collections of birds, mammals, plants, and herps relative to much of “mainland” China, although the warm temperate areas of Southern China appear well collected for some groups such as plants and herps making the absence of mammals and birds from this region even more striking. Much of the cold vastness of Russia too seems undersampled for natural history specimens. Yet bright spots on the Russian map shed light on the importance of local museums in the pursuit of cataloging biological diversity. For example, the high concentration of fungi samples near the western central part of the country is not a centroid artifact but the Yugra State University Biological Collection: an institution that has been collecting mycological specimens since the early 20th century [55]. Lack of digitization, collecting efforts, climate, and politics probably have a great deal to do with the lack of vouchered samples from the world’s largest country.

**Fig 4.**
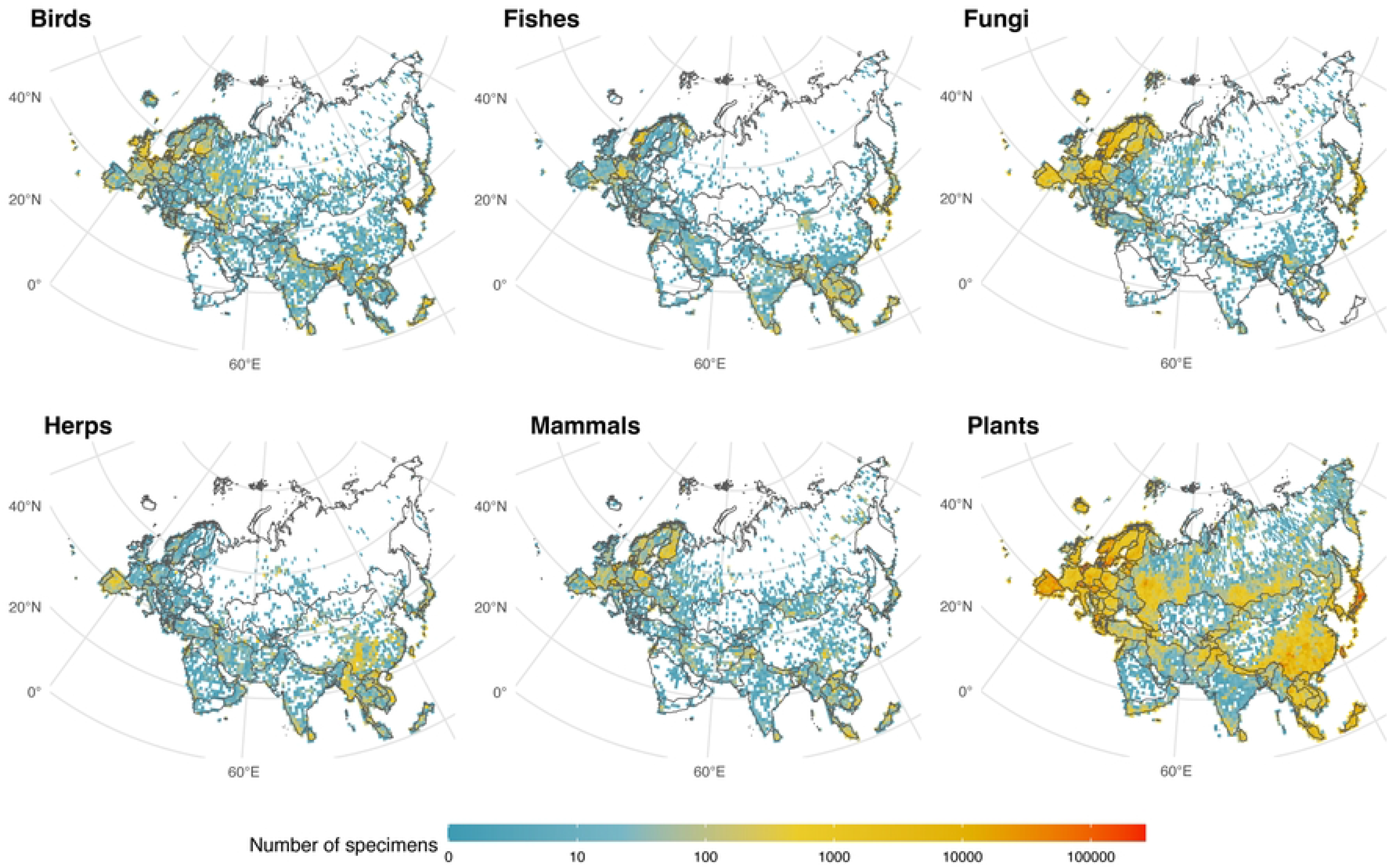
Maps of collecting efforts on Eurasia. Sample (voucher) quantity for each taxon considered in this study is shown.

**Fig 5.**
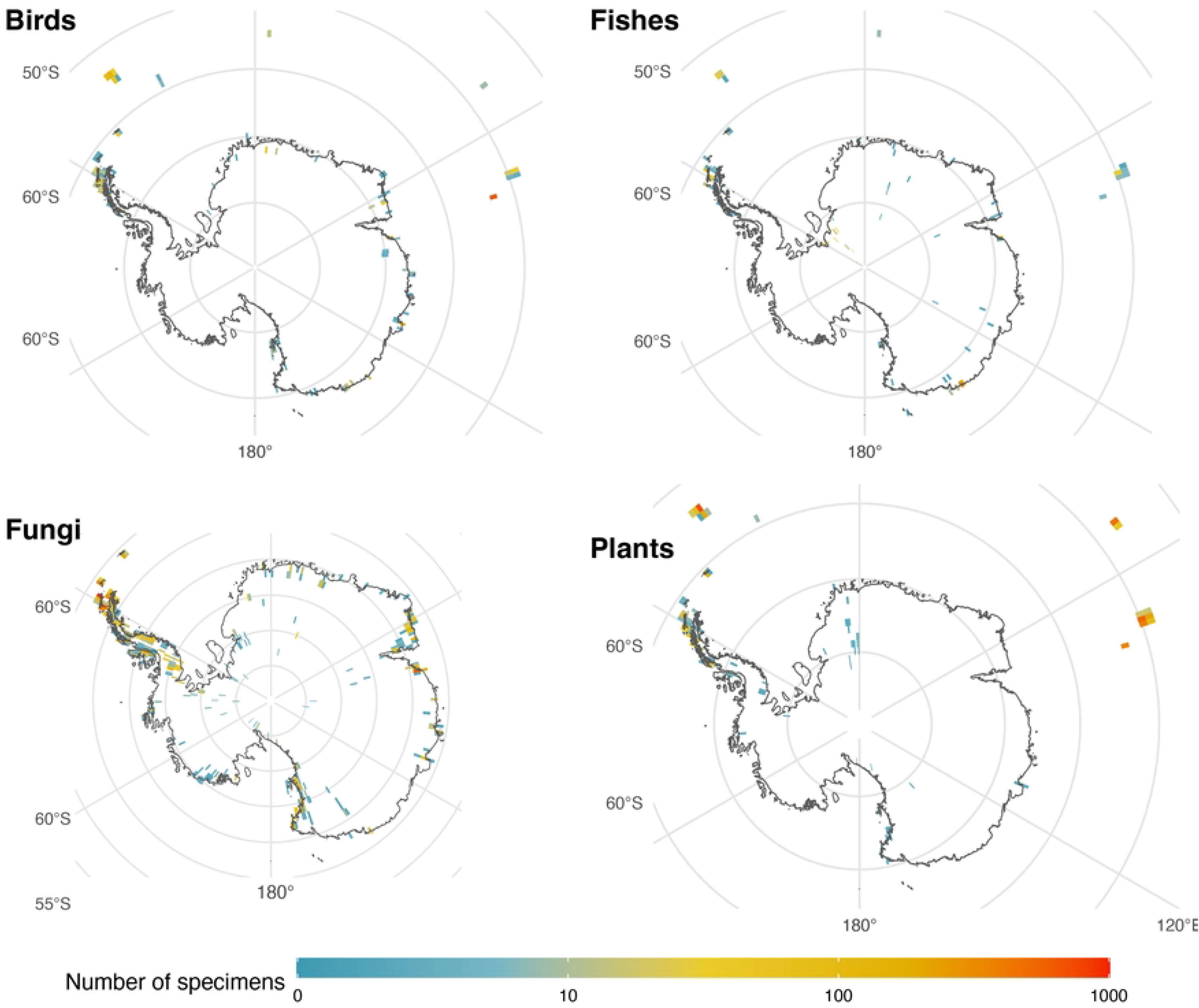
Maps of collecting efforts on Antarctica. Sample (voucher) quantity for each taxon considered in this study is shown.

### Antarctica

Antarctica is notable only in the absence of many collections for obvious reasons, there are no collections of freshwater fishes, amphibians or reptiles because none survive on the continent. Large mammal and bird collections do exist but are generally restricted to historic collections or from areas near the coasts including some not yet recorded on GBIF, similarly, lichen and plants are generally restricted to ice-free zones off of the continent (Fig 5). Observations of Antarctic fauna have been made but many of these are of invertebrates and fishes below the sea ice [56].

## Conclusions

The term “unexplored” can be used in many unrelated contexts, and it may not accurately reflect the state of knowledge in a given area. To some, “unexplored” may mean an area that has never been seen by human beings, while to others, it may mean an area that is not exploited by Westerners or is uninhabited. In examining the so-called “unexplored” places that remain on land, we find that a lack of digitized natural history collections may be a common cause for this label. In some cases, there may be a lack of natural history samples from an area because there are relatively few organisms there to collect (e.g., the Atacama Desert in Chile and Sahara Desert of Africa which are some of the driest places on Earth, as well as many parts of Antarctica). In some regions, political unrest has prevented access to naturalists (e.g., Libya, Venezuela, Colombia), and a lack of infrastructure for obtaining permits and permissions hurt both the global understanding of a region and local biodiversity education. Other areas appear simply to have been overlooked by natural history research— the limited number of practicing naturalists with even more limited funding may be an explanation for some of these regions remaining poorly researched. The term “unexplored” in these contexts can be harmful, as such terminology may inadvertently perpetuate misconceptions or undermine the value of existing research and the efforts of local scientists and indigenous people or disregard the underlying resource-imbalances impeding natural history research, collection, and digitization in many regions. For these reasons, we advocate for retiring the word “unexplored” for more precise and inclusive language (e.g., “this region lacks digitized natural history collections”) and suggest “biodiversity blindspots” as an alternative.

In this paper, we highlight areas lacking or having few collections, not to encourage their exploitation, but to call for an increased understanding of these areas in the context of global biological diversity for the sake of conservation and recognition in a natural history context. A multitude of factors have kept these places from the growing knowledge base of natural history and biodiversity, and in some cases, particularly where indigenous stakeholders are the protectors [57], traditional Western approaches to science may not be the most effective means of including these regions [58]. Good faith partnerships and collaborations, a fair exchange of knowledge and resources, and the acknowledgement that there are many different ways of knowing that are equally contributable, will be necessary starting points to conversations related to natural history in many of these areas (e.g., large swaths of the Amazon Rainforest that are home to uncontacted tribes). Recent calls for equity-based fieldwork should be heeded, and special care should be taken to avoid practicing parachute science, a term that refers to situations in which researchers, often from more economically developed or privileged countries, visit other marginalized countries to conduct research and then return to their home institutions without genuinely engaging or collaborating with locals [59–61]. Some of the most biologically diverse and well collected countries are also among the most exploited and experience high incidences of parachute science. Any future work in undercollected regions should be performed in ways that are considerate of and inclusive to local researchers and communities.

Accessibility to financial resources that make digitization (especially large-scale efforts) possible are a significant barrier for many museums and institutions— even for some larger institutions in the West. (e.g., based on GBIF data most natural history specimen collections from Costa Rica, Mozambique, and India are found outside of those countries). There are large museums that are not yet digitized and part of GBIF in some of the most biodiverse places, including but not limited to the Museum of Zoology of the University of São Paulo, the Museu Paraense Emílio Goeldi, Inpa Coleçoes Zoológicas, and the National Museum in Rio de Janeiro (which suffered a devastating fire in 2018; [62]). Modernizing access to the reference vouchers housed at these institutions would likely benefit research groups globally and facilitate natural history and conservation research. However, international digitization efforts will require sharing resources to create scientific infrastructure where it is lacking, especially in places where biodiversity is richest and the people poorest. It should be noted that while there are many benefits to digitization, care needs to be taken to not perpetuate colonialism. The benefits of digitization will not be realized equitably without global collaborative efforts that are inclusive to regions with fewer economic resources, and offer a fair exchange knowledge, technology, and support to empower these areas in their scientific endeavors.

The goal of this study was to increase our knowledge of “biodiversity blindspots” by highlighting these poorly-collected or undigitized places, and we hope these focus areas can be topics of discussion by researchers, politicians, indigenous stakeholders, and others who are interested in the discovery-based science of biodiversity research.

## Materials and Methods

Occurrence records of preserved specimens with GPS coordinates were downloaded from GBIF between July 1 and December 31, 2022 (Global Biodiversity Information Facility; https://www.gbif.org/) for each of six taxonomic groups: fungi, terrestrial and freshwater vascular plants, freshwater fishes, herpetofauna (amphibians and reptiles), birds, and mammals. As we are interested in general trends in collection records across continents, only occurrence records with associated voucher specimens were considered for each of the six taxonomic groups included in this study (we did not include observational data where no specimens were collected). To maximize the amount of georeferenced data retrieved from GBIF, we did not apply any other filters (e.g., continent, year) as georeferenced records may be missing information for such fields. Given the size of the vascular plant’s dataset (>126 million records), we split records by continent in R v.x.x (readr package; [63]) using the maximum and minimum latitude and longitude for each continent.

Available records from GBIF for each dataset (taxonomic group and data subsets for vascular plants) were cleaned in QGIS v.3.10.9 (QGIS.org, 2023. QGIS Geographic Information System. QGIS Association. http://www.qgis.org), to include records that fall within continent limits (including islands) using the “World Countries” shapefile as downloaded from ArcGIS. We used QGIS to split the world map into separate shape files corresponding to six regions: South America (with adjacent islands), North America (including landmasses in the Gulf of Mexico, Caribbean and west to Hawaii), Africa (including Madagascar and adjacent landmasses), Australia + Oceania (including islands in Oceania such as the Indonesian archipelago and New Zealand), Eurasia (including all of continental Europe and Asia as well as adjacent islands such as the British Isles, Iceland, Svalbard Archipelago, Japan and the Philippines), and Antarctica. These continent shape files were used to define the extent of polygon grids, which were in turn used to count the number of occurrence records per one-degree squared grid cell. Occurrence records that did not intersect with continent layers were removed. This procedure also removed data points exactly at 0°, 90°, or ≥180°, or in the ocean. Using the “count points in polygon” tool, the number of collections per grid cell were calculated across each continent. The number of collections in grids were log10 transformed using Field Calculator in QGIS. Collection gaps were then visually identified. To assess whether global differences in collection intensity could be due to climatic differences (e.g., lower species diversity in hot desert areas might lead to fewer collections in these regions - see [64]), collections maps were also compared with Köppen-Geiger climate maps (http://koeppen-geiger.vu-wien.ac.at/present.htm).

An interactive map was generated using QGIS2Web plugin (ver. 3.16.0) and can be found at https://canon-network.github.io/ under the “Specimen Database” tab. This map shows all collected specimen numbers divided into one-degree squared grid cells covering the globe and can be toggled for individual taxonomic groups. In the static maps shown below (Figs 1-5) lighter areas are poorly sampled regions based on GBIF records, while darkly colored (red) regions are well-collected (100K+ specimens) with many specimens from those areas. The specimen data was log-transformed to increase the color contrast of poorly sampled areas so they would be pronounced well on the maps. Non-log transformed data can be visualized in the interactive map described earlier in this paragraph.

## Acknowledgments

Greg Thom and Nathan Lord helped at various stages of manuscript preparations. We thank the thousands of natural history researchers, explorers, and collectors who contributed to the samples we examined here. We also thank the hundreds of museum researchers who prepared, digitized, and curated these voucher samples. We are also grateful to the databasing experts, particularly those at the Global Biodiversity Information Facility who have made this type of information widely accessible.

